# Parvalbumin interneuron mGlu_5_ receptors govern sex differences in prefrontal cortex physiology and binge drinking

**DOI:** 10.1101/2023.11.20.567903

**Authors:** Carly B. Fabian, Nilah D. Jordan, Rebecca H. Cole, Lily G. Carley, Shannon M. Thompson, Marianne L. Seney, Max E. Joffe

**Affiliations:** Department of Psychiatry, University of Pittsburgh, Pittsburgh, PA, 15219, USA; Translational Neuroscience Program, University of Pittsburgh, Pittsburgh, PA; Center for Neuroscience University of Pittsburgh, Pittsburgh, PA

**Keywords:** prefrontal cortex, alcohol, G protein-coupled receptor (GPCR), cognition, GABA, addiction

## Abstract

Despite established sex differences in the prevalence and presentation of psychiatric disorders, little is known about the cellular and synaptic mechanisms that guide these differences under basal conditions. Proper function of the prefrontal cortex (PFC) is essential for the top-down regulation of motivated behaviors. Activity of the PFC is tightly controlled by parvalbumin-expressing interneurons (PV-INs), a key subpopulation of fast-spiking GABAergic cells that regulate cortical excitability through direct innervations onto the perisomatic regions of nearby pyramidal cells. Recent rodent studies have identified notable sex differences in PV-IN activity and adaptations to experiences such as binge drinking. Here, we investigated the cellular and molecular mechanisms that underlie sex-specific regulation of PFC PV-IN function. Using whole-cell patch clamp electrophysiology and selective pharmacology, we report that PV-INs from female mice are more excitable than those from males. Moreover, we find that mGlu_1_ and mGlu_5_ metabotropic glutamate receptors regulate cell excitability, excitatory drive, and endocannabinoid signaling at PFC PV-INs in a sex-dependent manner. Genetic deletion of mGlu_5_ receptors from PV-expressing cells abrogates all sex differences observed in PV-IN membrane and synaptic physiology. Lastly, we report that female, but not male, PV-mGlu_5_^-/-^ mice exhibit decreased voluntary drinking on an intermittent access schedule, which could be related to changes in ethanol’s stimulant properties. Importantly, these studies identify mGlu_1_ and mGlu_5_ receptors as candidate signaling molecules involved in sex differences in PV-IN activity and behaviors relevant for alcohol use.

## Introduction

Sex differences contribute to the incidence, presentation, and severity of several psychiatric disorders [1,2]. For instance, women with major depressive disorder (MDD) experience greater symptom severity [3,4], men have an earlier age of onset of schizophrenia [5–7], and women display a faster progression through disease milestones of alcohol use disorder (AUD) [8,9]. Together, the broad sex differences in psychiatric diseases motivate continued preclinical research to identify the cells and plasticity mechanisms that mediate sex differences in brain function.

The prefrontal cortex (PFC) provides top-down regulation of motivated behaviors and displays sex differences in form and function [10–14]. Most PFC neurons are glutamatergic pyramidal cells that send, receive, and integrate information to and from various subcortical and cortical structures [15,16]. Pyramidal cell activity is tightly regulated by a small number of inhibitory neurons that transmit γ-aminobutyric acid (GABA). Parvalbumin-expressing inhibitory neurons (PV-INs) form a key subpopulation of these GABAergic cells which synapse onto perisomatic regions of pyramidal neurons and display fast-spiking firing properties [17–19]. Dysregulation of PFC PV-IN activity has been implicated in the etiology of many psychiatric disorders [20–23]. A growing body of literature in rodent models has also identified notable sex differences in PV-IN activity and adaptations to experiences. Indeed, male and female mice display sex differences in developmental trajectory of PFC PV expression [24], stress sex-specifically alters PFC PV-IN function [25–27], and PFC PV-INs undergo sex-dependent adaptations following four-weeks binge drinking [28]. Taken together, these studies suggest that sex differences in molecular mechanisms governing synaptic plasticity and intrinsic properties on PV-INs are critical in mediating experience-dependent adaptations to PFC function.

The group 1 metabotropic glutamate (mGlu) receptors, mGlu_1_ and mGlu_5_, are expressed on PFC PV-INs [29–32] and regulate cellular processes related to membrane physiology, synaptic strength, and endocannabinoid plasticity [33–35]. In addition, there are sex differences in signaling and function of mGlu_1_ and mGlu_5_ receptors [36–38]. Here, we tested the hypothesis that mGlu_1_ and mGlu_5_ receptors mediate sex differences in PFC PV-IN function. Using genetically engineered mice, whole-cell patch clamp electrophysiology, and selective pharmacology, we examined sex differences in PV-IN function and regulation by mGlu_1_ and mGlu_5_ receptors. We then assessed a causal role for mGlu receptors in regulating PFC PV-IN function through mice with cell type-specific deletion of mGlu_5_ receptors. The present results support the hypothesis that PV-IN mGlu_5_ receptor signaling is critical for sex differences in PFC physiology and drinking.

## Materials and Methods

### Mice

Mice were maintained on a 12:12 light-dark cycle (lights on at 6:00 AM) with food and water *ad libitum*. Mice were bred at Charles River Laboratory (Wilmington, MA) and delivered to the University of Pittsburgh at 6-8 weeks of age. All mice were on a congenic C57BL/6J background and experiments began in adulthood (>10 weeks). Sex determination was limited to an assessment of external genitalia. Mice expressing tdTomato in PV-INs were bred by crossing female homozygous PV-Cre mice [39] (Jackson Laboratories, 017320) with male homozygous Rosa26-loxP-STOP-loxP-CAG-tdTomato “Ai9” mice [40] (Jackson Laboratories, 007909). PV-mGlu_5_^-/-^ mice were generated by crossing PV-Cre and Ai9 mice with those harboring a floxed *Grm5* locus [31,41] (Jackson Laboratories, 028626). All experiments were approved by the University of Pittsburgh IACUC and conducted in accordance with the NIH Guidelines.

### Brain slice preparation

Mice (N=68) were decapitated under isoflurane anesthesia; brains were immediately dissected and placed in *N*-methyl-D-glucamine (NMDG) cutting solution, (in mM: 93 NMDG, 20 HEPES, 2.5 KCl, 0.5 CaCl_2_, 10 MgCl_2_, 1.2 NaH_2_PO_4_, 25 glucose, 5 Na-ascorbate, and 3 Na-pyruvate). Coronal slices (300-μM) of the PL-PFC were prepared in room-temperature NMDG solution. Slices recovered for 10 minutes in warm (32-35°C) NMDG solution and were then transferred to room temperature artificial cerebral spinal fluid (ACSF) (in mM:119 NaCl, 2.5 KCl, 2 CaCl_2_, 1 MgCl_2_, 1 NaH_2_PO_4_, 11 glucose, and 26 NaHCO_3_) for at least 1 hour before recordings. All solutions were oxygenated (95% O_2_ / 5% CO_2_).

### Ex vivo whole-cell patch-clamp electrophysiology

Data were acquired using a Multiclamp 700B amplifier and pCLAMP 11 software (Molecular Devices). Slices were transferred to the recording chamber and perfused at 2mL/min with heated ACSF (30-32°C). For all whole-cell patch-clamp recordings, cells were patched with a borosilicate glass pipette pulled to 3-5 MΩ with a horizontal electrode puller (P-1000, Stutter Instruments). Pipettes were filled with a potassium-based internal solution (in mM: 125 K-gluconate, 4 NaCl, 10 HEPES, 0.1 EGTA, 4 MgATP, 0.3 Na_2_GTP, 10 Tris-phosphocreatine). PV-INs in layer 5 of the PL-PFC were identified by tdTomato fluorescence and cell identity was confirmed via characteristic fast-spiking phenotype. After gaining whole-cell access, PV-INs were dialyzed for 5 minutes under voltage-clamp at -80mV. Membrane properties were then assessed under current-clamp as described [42]. A series of current injections (1s, starting at - 150pA, ending at +500pA, increments of 25pA) were applied to patched cells. Resting membrane potential (V_m_) was determined as the average baseline membrane potential prior to current injections. Membrane resistance (R_m_) was calculated as the slope of the hyperpolarizing membrane potential over three injections [-75, -50, -25pA]. Rheobase was determined as the minimal current injection needed to elicit firing.

Cells were then returned to voltage-clamp configuration. Spontaneous excitatory postsynaptic currents (sEPSCs) were electrically isolated at -80mV, near the reversal potential for chloride [28]. DHPG-induced changes in depolarizing currents and excitatory transmission onto PFC PV-INs were assessed sequentially in the same cell. Following a 5-minute baseline, PV-INs underwent sequential 5-min application of 5μM and then 50μM DHPG. In some experiments, LY367385 (100μM) or MTEP (3μM) was applied prior to baseline recordings and was present for the entire experiment. The holding current did not drift in 15-minute vehicle control experiments (0.3 ± 3.5pA, n=5 cells). AMPAR-mediated sEPSCs on PV-INs were identified using pattern-matching event detection software (ClampFit). The template was generated from PV-IN recordings (match threshold 3.5) and did not detect events in NBQX. Change in holding current and sEPSC frequency were calculated by comparing the average values over the last 2 minutes of drug application to the last 2 minutes of baseline. At the conclusion of the 5-minute 50µM DHPG application, cells were returned to current-clamp and membrane properties were re-assessed.

In separate PV-INs, endocannabinoid-mediated short-term plasticity was assessed via depolarization-induced suppression of excitation (DSE) [43,44]. Bipolar nichrome wire stimulating electrodes were positioned in layer 2/3 medial to the recording electrode. EPSCs were evoked with a single pulse (10-50μA) at 0.2Hz to elicit EPSCs between 100-200pA. After establishing a stable 1-minute baseline, DSE was elicited by depolarizing patched PV-INs to +20mV for 10 seconds without stimulation, followed by an immediate return to -80mV and electrical stimulation. DSE was repeated 3-4 times per cell under basal conditions. DHPG (5μM) was then applied for 5 minutes and the DSE protocol was repeated. Changes in the EPSC amplitude were normalized to baseline. To assess inhibitory transmission, some PV-INs were patched with a high-chloride cesium-based internal solution (in mM: 125 CsCl, 5 NaCl, 10 HEPES, 4 MgATP, 0.3 Na_2_GTP, 10 Tris-phosphocreatine, 5 Qx-314-Br) and held at -70mV in NBQX (10 µM). Data for electrophysiology studies was processed using ClampFit (Molecular Devices).

### Intermittent access (IA) ethanol

IA ethanol was performed as described [42,46]. For three 24-hour periods per week, ethanol was provided in home cages, with 24-48 hours between each exposure. Ethanol was diluted from undenatured 95% ethyl alcohol and was delivered by modified conical tubes with rubber stoppers and open tip tubes. Ethanol was generally provided and removed 3-4 hours prior to start of the dark cycle. After a ramp during the first three sessions [3, 6, 10%], the ethanol concentration was set at 20% ethanol for the remainder of the study.

### Locomotor response to ethanol

We assessed the locomotor response to ethanol in mice without previous exposure to ethanol. Mice received saline vehicle (10 µL/kg, *i.p.*) or ethanol (0.5, 1, or 2 g/kg) immediately before being placed in an open field chamber for 20 minutes. Over 4 consecutive days, all mice received all doses in a counterbalanced Latin-square design (see schematic in Figure 5). Location and distance traveled were determined by video-based tracking (ANY-maze, Stoelting).

### Drugs

Drugs were purchased from HelloBio. Concentrations were based on previous studies [29,32,47,48]. DHPG has no effect on Group 2 or Group 3 mGlu receptors (mGlu_2,3,4,7,8_) at concentrations up to 1mM [49,50]. Drugs were formulated, aliquoted, and frozen at 1000X, then thawed and diluted in ACSF on the day of recording. LY367385 [(*S*)-(+)-α-Amino-4-carboxy-2-methylbenzeneaceticacid] stock was made in 0.1M NaOH. MTEP [3-((2-Methyl-1,3-thiazol-4-yl)ethynyl)pyridine hydrochloride], DHPG [(*RS*)-3,5-Dihydroxyphenylglycine], and NBQX stocks were made in water.

### Statistics

Data are presented as mean±SEM or box plots (min, Q1, median, Q3, max). Mean±SEM for key experiments is reported in Table S1. Statistical analyses were performed using GraphPad Prism. For electrophysiology, *n* represents the number of cells. For all analyses, p<0.05 was considered significant and p<0.07 was labeled trending. Data were analyzed using two-tailed Student’s t-test. Two-way or three-way ANOVA, or mixed-effects analysis, with Bonferonni post-hoc comparisons were used for analyses requiring multiple comparisons. Statistical analyses are reported in the legends and in Table S2.

## Results

### PL-PFC PV-INs from female are mice are more excitable than PV-INs from males

To assess intrinsic physiology of PFC PV-INs, we made whole-cell patch-clamp recordings from genetically engineered mice expressing tdTomato in PV-positive cells [32,42,51]. All (n=75/75) tdTomato-expressing cells from layer 5 of the PL-PFC displayed characteristic fast-spiking interneuron properties [52,53], including: high frequency firing, low input resistance, and minimal spike-frequency adaptation (Figure 1A). In general, PV-INs from male and female mice displayed comparable basal membrane properties, without significant differences in resting potential, membrane resistance, or medium afterhyperpolarization (Figure S1A-C). However, in response to positive current injections, cells from female mice exhibited greater spike-firing compared to cells from males (Figure 1B) and fired action potentials in response to a lower threshold current (Figure 1C). Together, these data indicate that while PV-INs exhibit similar basal membrane properties between sexes, PV-INs from female mice are more excitable than those from males.

**Figure 1.**
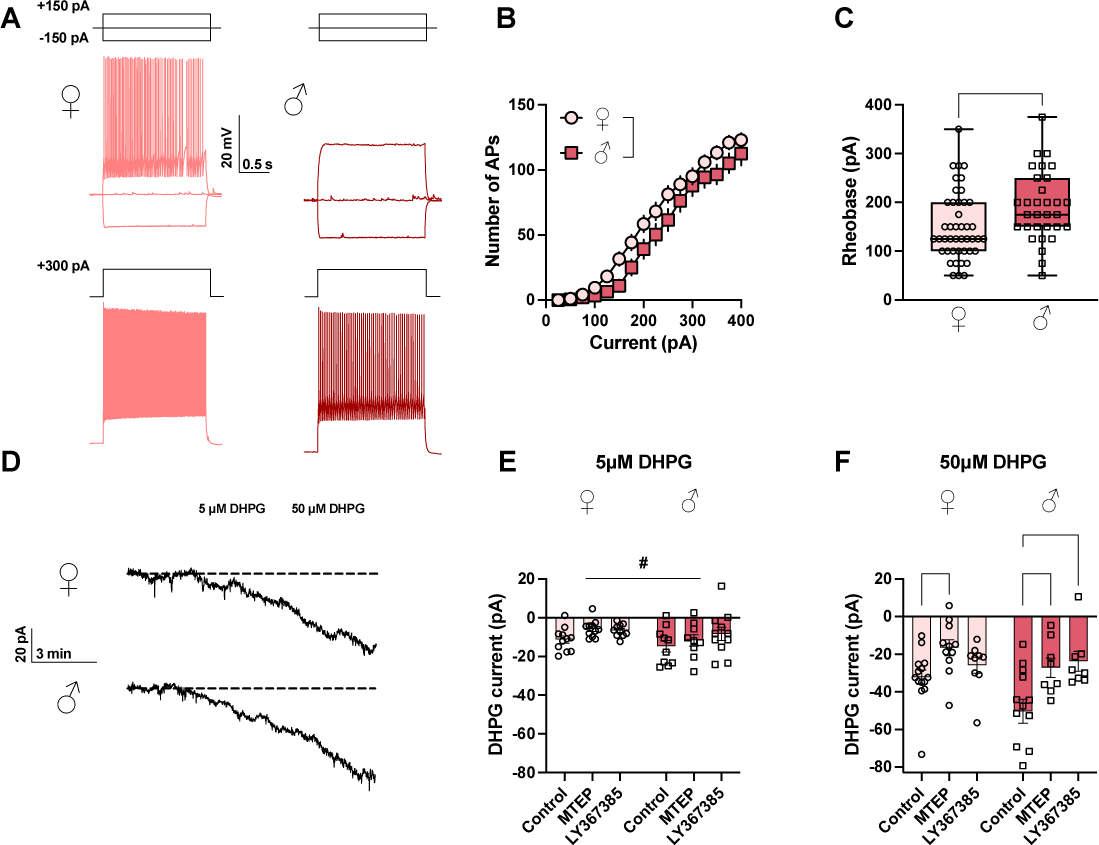
PV-INs from female mice are more excitable at baseline, and PV-INs from male mice are more responsive to mGlu_1/5_ receptor stimulation. **(a)** Representative PV-IN current-clamp recordings from female (left, pink) and male (right, red) control mice. Traces depict responses to step current injections. The responses to -150 pA, 0 pA, +150 pA are shown on top and the responses to +300 pA are shown on the bottom. Scale bars 20mV, 0.5s. (**b**) PV-INs from female mice display greater action potential (AP) frequency relative to males in response to step current injections (Two-way mixed-effects analysis, main effect of sex: F_1,73_ = 4.1, *: p<0.05; sex x current interaction: F_15,1005_ = 2.0, p<0.05). n/N = 33-42/15-19 cells/mice per group. (**c**) PV-INs from female mice display lower rheobase than males (t_73_ = 2.5; *: p<0.05). (**d**) Representative traces showing effects of 5 μM and 50 μM DHPG on holding current in PV-INs from female (top) and male (bottom) mice. Scale bars indicate 20 pA, 3 min. (**e**) At a threshold concentration (5 μM), DHPG induced a trend toward greater depolarizing currents in male PV-INs than females (Two-way ANOVA, main effect of sex: F_1,56_ = 3.4, #: p<0.072; main effect of antagonist: F_2,56_ = 2.9, p<0.065). n/N = 9-11/8-10 cells/mice per group. (**f**) A saturating concentration (50 μM) of DHPG induced greater depolarizing inward currents in PV-INs from male mice. MTEP attenuated the effects of 50 μM DHPG in both sexes. In male mice only, LY367385 blocked DHPG-induced depolarizing currents in PV-INs. (Two-way ANOVA, main effect of sex: F_1,57_ = 4.3, p<0.05; main effect of antagonist: F_2,57_ = 9.3, p<0.001; *p<0.05, **: p<0.01 Bonferroni post-tests). n/N = 8-14/8-10 cells/mice per group.

### mGlu_1_ and mGlu_5_ receptors depolarize PFC PV-IN in a sex-dependent manner

We and others previously demonstrated that mGlu_1_ and mGlu_5_ receptors modulate PV-INs in frontal cortex [29–31], but most of these studies were limited to males. We therefore aimed to determine whether mGlu_1_ and mGlu_5_ receptors modulate PV-IN function in a similar manner in males and females. We examined sensitivity to mGlu_1_ and mGlu_5_ receptor stimulation using a threshold concentration of the non-selective mGlu_1/5_ agonist DHPG (5μM) alone, in the presence of the selective mGlu_5_ NAM MTEP (3μM), or in the presence of the selective mGlu_1_ antagonist LY367385 (100μM) (Figure 1D). Threshold DHPG induced modest depolarizing currents in PV-INs of both sexes; however, PV-INs from male mice displayed a trend towards greater depolarizing changes in holding current compared to females (Figure 1E), with no effect of either antagonist. We next assessed a high concentration of DHPG (50μM), which induced greater depolarizing current relative to the threshold concentration in both sexes (Figure 1F). Depolarizing currents in response to 50μM DHPG were greater in PV-INs from males, and selective pharmacological antagonism revealed additional sex differences in the receptor subtypes involved. In PV-INs from all mice, MTEP attenuated DHPG-induced depolarizing currents. By contrast, LY367385 reduced DHPG currents in PV-INs from male mice but did reach significance in females. Dual application of both MTEP and LY367385 attenuated but did not completely block DHPG currents in both male and female PFC PV-INs (Figure S2A), potentially due to competition between DHPG and LY367385 at the orthosteric site or partial/biased NAM activity of MTEP in this assay. Together, these data suggest that PFC PV-IN membrane physiology is modulated by mGlu_5_ receptors across sexes, but that PV-INs from male mice are more sensitive to mGlu_1_ receptor activation. We also returned cells to current-clamp at the conclusion of the DHPG wash-on and found PV-INs from males were depolarized to a greater extent than females (Figure S3A).

### mGlu_1_ and mGlu_5_ regulate synaptic strength on PFC PV-INs in a sex-dependent manner

In addition to membrane physiology, Group 1 mGlu receptors can modulate synaptic strength on many cell types, including other types of interneurons in frontal cortex [32,54–57]. We measured AMPA receptor-mediated sEPSCs under voltage-clamp and found no evidence for basal sex differences in excitatory or inhibitory synaptic strength (Figure S1D-I). We next determined whether Group 1 mGlu receptors regulate excitatory transmission. Threshold DHPG (5μM) had no effect on sEPSC frequency (Figure 2A and 2B), but a higher concentration of DHPG (50μM) induced a three-fold increase in sEPSC frequency in PV-INs across sexes (Figure 2C). Further pharmacological studies revealed a sex-specific receptor mechanism underlying this effect: LY367385 blocked enhancement of excitatory drive in PV-INs from male mice, while the effects of DHPG on sEPSC frequency were attenuated by MTEP in PV-INs from females. Dual application of MTEP/LY367385 blocked the DHPG-induced increase in excitatory drive (Figure S2B). These findings suggest a sex-specific role of the group 1 mGlu receptors in regulating synaptic strength onto PFC PV-INs, primarily driven by mGlu_1_ receptor signaling in males and by mGlu_5_ receptors in females.

**Figure 2.**
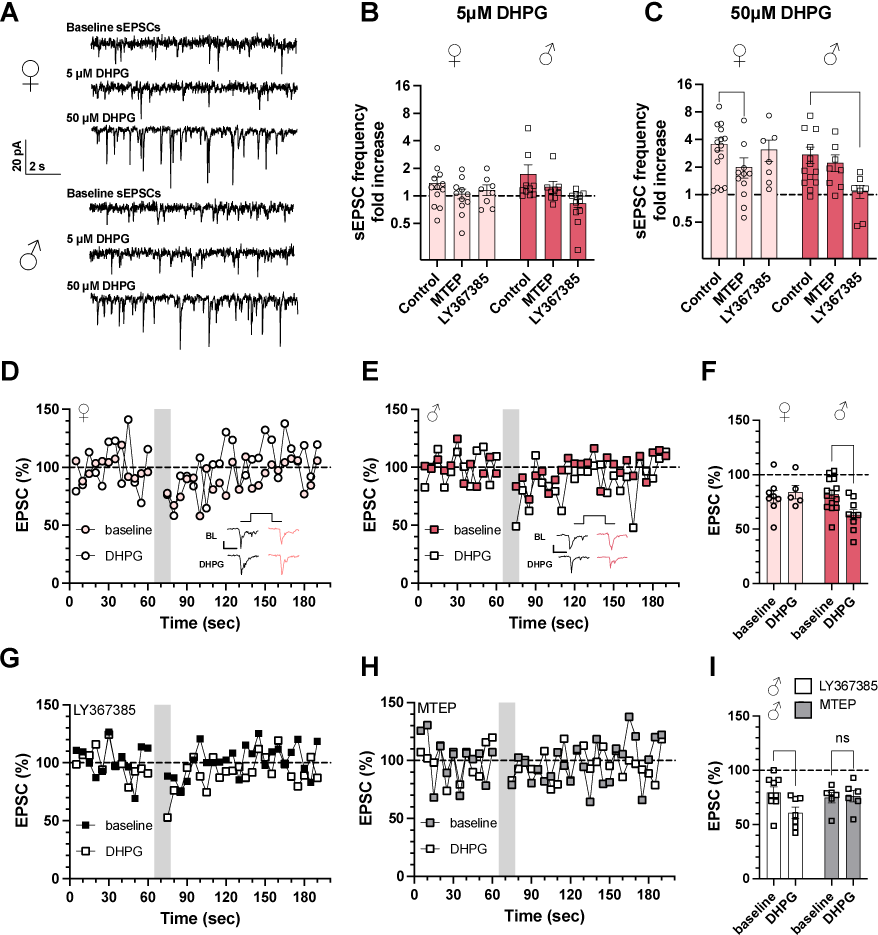
mGlu_1_ and mGlu_5_ receptors sex-specifically regulate synaptic strength and endocannabinoid plasticity in PV-INs. **(a)** Representative voltage-clamp recordings displaying excitatory postsynaptic current (sEPSC) traces from PV-INs from female (top) and male (bottom) mice. Recordings are from pre-drug baseline period (top) and following sequential 5-minute applications of 5 μM (middle) and 50 μM (bottom) DHPG. 20pA, 2s. **(b)** Minimal effects of 5 μM DHPG application on sEPSC frequency. LY367385 attenuated the effects of 5uM DHPG (Two-way ANOVA, main effect of antagonism: F_2, 56_ = 4.0, p<0.05). n/N = 8-12/8-10 cells/mice per group. **(c)** 50 μM DHPG induced a three-fold increase in sEPSC frequency in PV-INs from male and female mice. LY367385 inhibited DHPG-induced enhancement of sEPSC frequency in PV-INs from male mice, whereas MTEP attenuated this effect in PV-INs from female mice. (Two-way ANOVA, main effect of sex: F_1,56_ = 4.2, p<0.05; main effect of antagonism: F_2,56_ = 3.4, p<0.05; antagonism x sex interaction: F_2,56_ = 3.4, p<0.05; *: p<0.05 Bonferroni post-test). n/N =7-15/8-10. Differences in sEPSC frequency were assessed on log-transformed data. **(d)** Representative experiment of depolarization-induced suppression of excitation (DSE) elicited by depolarizing patched PV-INs from -80 mV to +20 mV for 10 seconds. DSE was averaged across three technical replicates at baseline, then again following application of threshold DHPG (5 μM). DHPG did not affect DSE in PV-INs from female mice. Representative traces of baseline EPSCs (black) and DSE (pink) prior to (top) and during 5 μM DHPG application (bottom). Scale bars indicate 200 pA, 20ms. **(e)** Representative DSE experiment in PV-IN from male mouse. **(f)** DHPG potentiated DSE in PV-IN from male but not female mice (Two-way mixed-effects analysis, DHPG x sex interaction: F_1,12_ = 5.5, p<0.05; ** p<0.01 Bonferroni post-test). Representative experiments of DSE in male PV-INs in the presence of **(g)** LY367385 and **(h)** MTEP. **(i)** DHPG potentiated DSE in the presence of LY367385 but the effect was blocked by MTEP (Two-way mixed-effects analysis, DHPG x antagonist interaction: F_1,24_ = 3.6, p<0.069; *: p<0.05 Bonferroni post-test). n/N = 5-14/5-8 cells/mice per group.

### mGlu_5_ receptors facilitate short-term plasticity at PV-INs from male but not female mice

We next sought to examine mechanisms of mGlu receptor synaptic plasticity. In several brain regions, activation of postsynaptic Group 1 mGlu receptors can trigger endocannabinoid release and reductions in neurotransmitter release [58–61]. We elicited endocannabinoid plasticity by depolarization-induced suppression of excitation (DSE) [62,63], a form of short-term depression mediated by CB1 receptor activation and reduced glutamate release probability. Under control conditions, PV-IN EPSC amplitude was reduced by approximately 20% by cell depolarization across sexes (Figure 2D and 2E). We next assessed whether mGlu_1_ and mGlu_5_ receptor activation can facilitate DSE by applying the threshold concentration of DHPG (5μM). While no effect was observed in female mice, threshold DHPG enhanced DSE in PV-INs from male mice (Figure 2E and 2F). Importantly, threshold DHPG did not affect EPSC amplitude in the absence of postsynaptic depolarization (Figure S4), suggesting the facilitation of DSE is a metaplastic change in endocannabinoid signaling, rather than an overall change in glutamate transmission. Based on our previous findings, we predicted that the LY367385 would block the enhancement of DSE in male mice. To our surprise, DHPG retained the ability to enhance DSE in LY367385 (Figure 2G and 2I), while MTEP completely blocked DSE potentiation (Figure 2H and 2I). We did not perform experiments using the antagonists in female mice because there was no effect of DHPG alone. Taken together, these findings indicate that mGlu_5_ receptors can potentiate endocannabinoid signaling in PV-INs from male mice.

### Genetic deletion of mGlu_5_ receptors from PV-INs abrogates sex-differences in membrane and synaptic physiology

We next used a genetic approach to examine mGlu_5_ receptor function within PV-INs. We bred mice to selectively ablate mGlu_5_ receptors from PV-expressing cells via Cre-mediated deletion of *Grm5* [31]. PV-mGlu_5_^-/-^ mice display postnatal reduction of mGlu_5_ receptors to ∼20% of control levels, with recombination beginning between around P28, reflecting endogenous PV expression. We first assessed PV-IN membrane properties in PV-mGlu_5_^-/-^ mice. In contrast to controls (Figure 1A-1C), we observed no sex differences in PV-IN spike-firing (Figure 3A and 3B) or rheobase (Figure 3C) in PV-mGlu_5_^-/-^ mice. Furthermore, we observed no sex differences in any other membrane property or in basal synaptic strength (Figure S5 and Table S2). These data suggest that mGlu_5_ receptors regulate the development and/or maintenance of sex differences in PV-IN excitability. We next assessed responses to mGlu_1/5_ receptor activation, hypothesizing that DHPG would not depolarize PV-INs in female PV-mGlu_5_^-/-^ mice. To our surprise, DHPG induced comparable depolarizing currents in both sexes (Figure 3D-3F), and currents were attenuated by either MTEP or LY367385. These findings suggest that (1) there is residual expression of mGlu_5_ receptors in PV-mGlu_5_ ^-/-^ mice, consistent with previous studies [31], and (2) female PV-mGlu_5_^-/-^ mice display unmasked sensitivity to mGlu_1_ receptor antagonism. We next assessed DHPG potentiation of excitatory drive in PV-mGlu_5_^-/-^ mice (Figure 4A). 5μM DHPG had little effect on sEPSC frequency (Figure 4B), while 50μM DHPG rapidly increased excitatory drive on PV-INs in both sexes (Figure 4C). In contrast to control mice, we observed a main effect of mGlu receptor antagonist without an effect of sex or interaction. Posthoc tests across sexes revealed that LY367385, but not MTEP, reduced the ability of DHPG to potentiate sEPSC frequency, again indicating loss of mGlu_5_ function in female mice and enhanced sensitivity to mGlu_1_ receptor antagonism. Finally, we performed DSE experiments to assess eCB signaling (Figure 4D-4F). DHPG failed to potentiate DSE in PV-INs of PV-mGlu_5_^-/-^ mice. Taken together, all sex differences in PV-IN physiology and mGlu receptor regulation were absent in PV-mGlu_5_^-/-^ mice.

**Figure 3.**
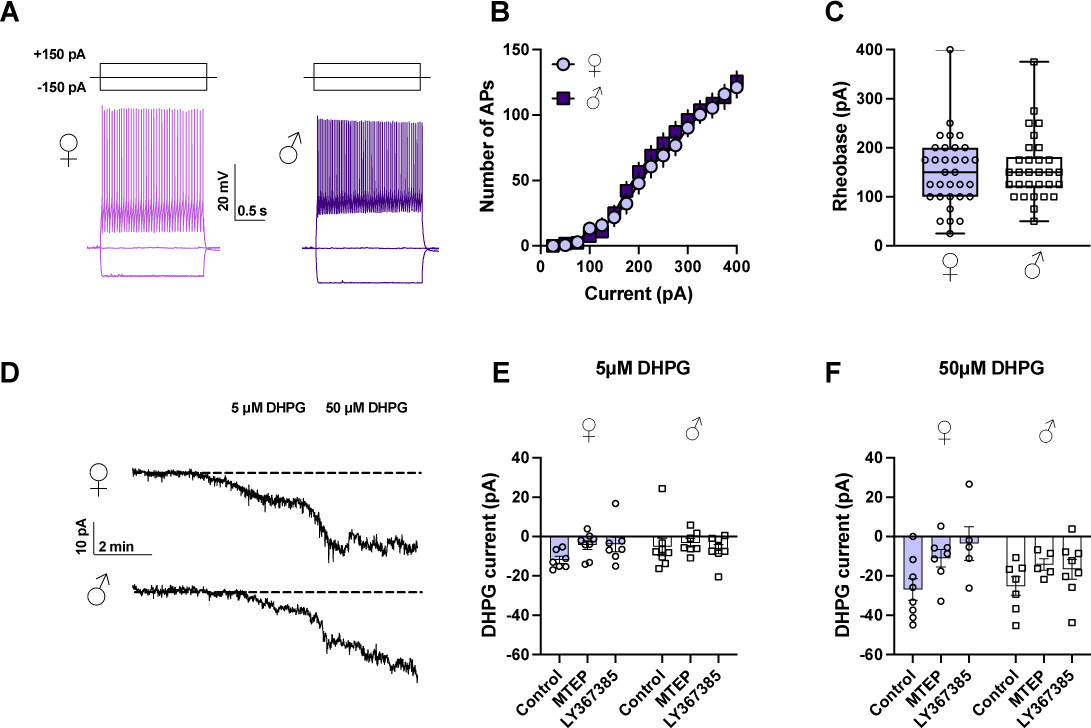
Cell type-specific genetic deletion of *Grm5* abrogates sex differences in PV-IN membrane physiology. **(a)** Representative PV-IN current-clamp recordings from female (left, light purple) and male (right, dark purple) PV-mGlu_5_^-/-^ mice. Traces depict responses to step current injections at -150 pA, 0 pA, +150 pA. **(b)** No differences in current-evoked spiking were observed in PV-INs from male and female PV-mGlu_5_^-/-^ mice. **(c)** PV-mGlu_5_^-/-^ mice display no sex differences in rheobase (t_60_ = 0.37, ns,). n/N= 30-32/5-8 cells/mice per group. **(d)** Representative traces showing effects of 5 μM and 50 μM DHPG on holding current in female (top) and male (bottom) PV-INs. Scale bars indicate 10 pA, 2 min. **(e)** No effect of sex or antagonist on depolarizing currents following threshold DHPG (5 μM). **(f)** Saturating DHPG (50 μM) induced depolarizing currents to a comparable magnitude in PV-INs from male and female mice. Across sexes, depolarizing currents were blocked by application of either MTEP or LY367385. (Two-way ANOVA, main effect of sex: F_1,34_ = 1.2, p=0.28; main effect of antagonist: F_2,34_ = 5.3, p<0.05, *: p<0.05 Bonferroni post-test). n/N = 5-9/5-8 cells/mice per group.

**Figure 4.**
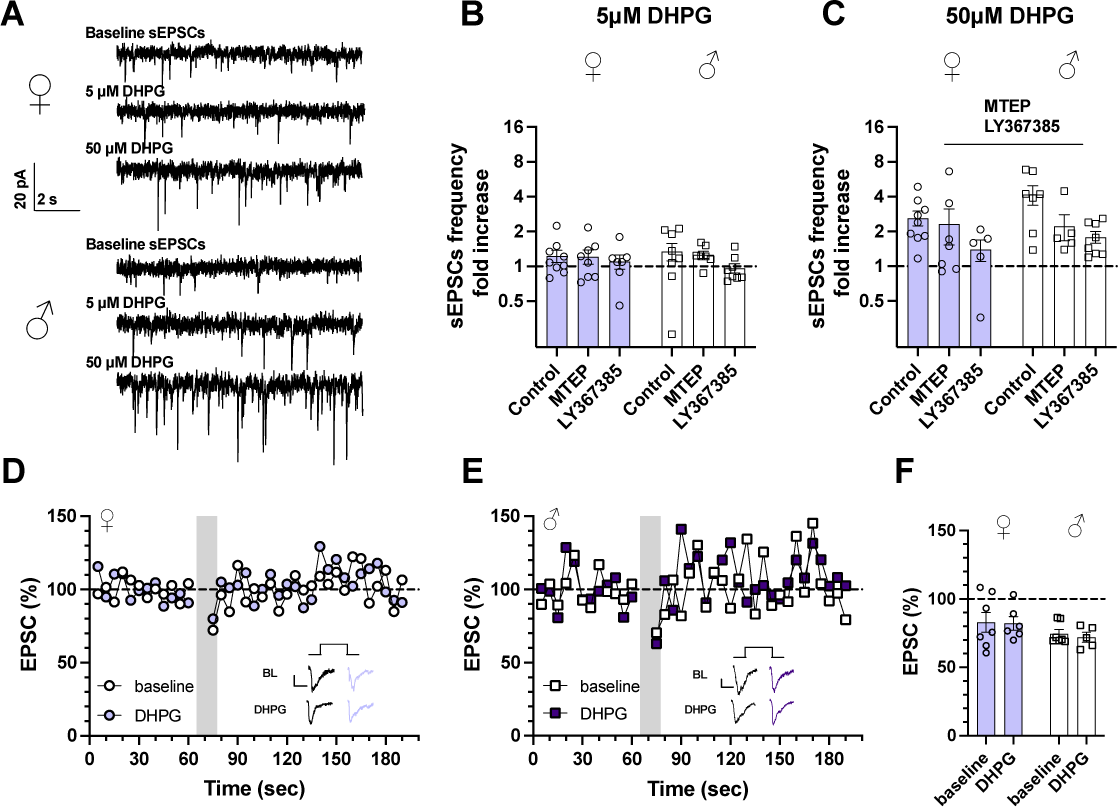
Cell-type specific genetic deletion of *Grm5* abrogates sex differences in PV-IN synaptic physiology. **(a)** Representative voltage-clamp recordings displaying sEPSC traces from PV-INs from female (top) and male (bottom) PV-mGlu_5_^-/-^ mice. Recordings are from pre-drug baseline period (top) and following sequential 5-minute applications of 5 μM (middle) and 50 μM (bottom) DHPG. Scale bars 20 pA, 2 s. **(b)** Minimal effects of 5 μM DHPG application on sEPSC frequency. **(c)** No sex differences in the effects of 50 μM DHPG on sEPSC frequency in PV-INs from PV-mGlu_5_^-/-^ mice, but LY367385 blocked the enhancement of excitatory drive across sexes. (Two-way ANOVA, main effect of sex: F_1,34_ = 2.8, p=0.099; main effect of antagonist: F_2,34_ = 6.0, p<0.01, *: p<0.05, **: p<0.01 Bonferroni post-test). Differences in sEPSC frequency were assessed on log-transformed data. n/N = 5-9/5-8 cells/mice per group. Representative experiments of DSE from **(d)** female and **(e)** male PV-mGlu_5_^-/-^ mice. Representative traces of baseline EPSCs (black) and DSE (pink) prior to (top) and during 5 μM DHPG application (bottom). **(f)** DHPG did not affect DSE in PV-mGlu_5_^-/-^ mice. n/N= 7-8/3-4 cells/mice per group.

### Deleting mGlu_5_ receptors from PV-INs in female mice, but not males, decreases voluntary drinking

mGlu_5_ receptor signaling regulates behaviors related to alcohol use [33,64], and previous studies from our group have demonstrated that drinking and ethanol reward learning alter the PV-IN physiology [28,51]. Based on this literature, and our current findings that mGlu_5_ receptors regulate synaptic strength onto PV-INs in female mice but not males, we hypothesized that mGlu_5_ receptors in PV-INs regulate binge drinking in a sex-dependent manner. To test this, we provided control and PV-mGlu_5_^-/-^ mice with two-bottle choice IA ethanol over 7 weeks (Figure 5A) [42,46]. As observed by our lab and many others [42,65–67], female mice drank more than male mice overall. Moreover, a sex by genotype interaction emerged, such that female but not male PV-mGlu_5_^-/-^ mice drank less than their respective controls (Figure 5B). Female mice had greater ethanol preference than males but there was no effect of genotype on this measurement (Figure 5C). We next performed follow-up studies in a new cohort of mice to assess sensitivity to ethanol’s stimulant properties (Figure 5D). Low-medium doses of ethanol increased locomotor activity. PV-mGlu_5_^-/-^ mice displayed enhanced hyperactivity relative to controls, but no effect of sex or interaction was observed. We assessed the time course of locomotor activity following the top dose (2 g/kg) and found that PV-mGlu_5_^-/-^ mice displayed increased locomotor activity during the first half of the test (Figure 5E). Finally, we found no differences in center time (Figure 5F), suggesting minimal effect of genotype on anxiety-like behavior. Together, these data implicate mGlu_5_ receptors in PV cells as a key mediator of sex differences in drinking, potentially related in part to ethanol’s stimulant properties.

**Figure 5.**
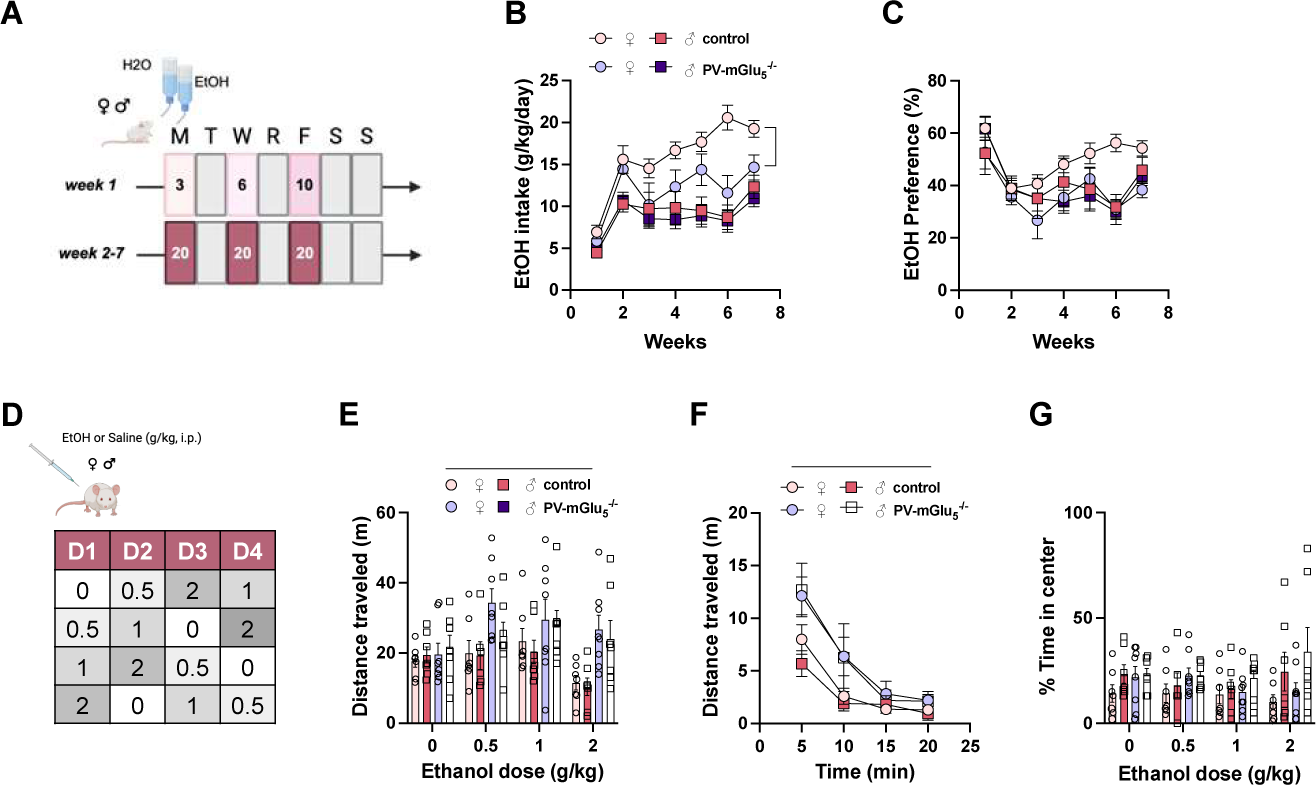
Female but not male PV-mGlu_5_^-/-^ mice display reduced binge drinking. **(a)** Schematic of intermittent access (IA) ethanol procedure. Mice had access to ethanol in a two-bottle choice procedure every other day for 7 weeks. **(b)** Female mice drank more than male mice overall (Three-way mixed-effects analysis, main effect of sex: F_1,34_ = 22.9, p<0.0001). Follow-up tests indicated that female PV-mGlu_5_^-/-^ mice drank less than controls (Two-way mixed-effects analysis, main effect of genotype: F_1,16_ = 7.5, p<0.05; genotype x sex interaction: F_6,93_ = 2.6, p<0.05), but no effect of PV-mGlu_5_^-/-^ was detected in male mice. N = 6-12 mice per group. **(c)** Female mice had a greater preference for ethanol more than male mice overall (Three-way mixed-effects analysis, concentration x sex interaction: F_6,199_ = 3.1, p<0.01), but there was no effect of genotype on ethanol preference. **(d)** PV-mGlu_5_^-/-^ mice displayed enhanced hyperactivity following ethanol injections, without any effect of sex (Three-way mixed-effects analysis, main effect of dose: F_3,75_ = 3.6, p<0.05; main effect of genotype: F_1,99_ = 19.3, ****:p<0.0001). N = 7-8 mice per group. main effect of genotype: F_1,99_ = 19.3, ****:p<0.0001). **(e)** PV-mGlu_5_^-/-^ mice displayed enhanced hyperactivity immediately after the 2 g/kg ethanol injection (Three-way mixed-effects analysis, main effect of genotype: F_1,25_ = 6.6, p<0.05; genotype x time interaction: F_3,75_ = 5.0, p<0.01). **(f)** Male mice spent more time in the center of the open field but no effect of sex or ethanol were detected (Three-way mixed-effects analysis, main effect of sex: F_1,25_ = 6.8, p<0.05).

## Discussion

Recent studies have identified sex differences in PFC PV-IN activity and adaptations to experiences under basal conditions and in disease models [25–27]. Understanding the molecular mechanisms mediating such sex differences in PV-IN function can help direct the development of novel therapeutic targets.

Here, we found that PV-INs from females exhibited lower rheobase and greater current-evoked spike-firing, while passive membrane properties, such as resting membrane potential and membrane resistance, did not differ between sex. These findings are consistent with prior studies showing modest sex differences in PFC PV-IN excitability, whereby PV-INs from female mice are more excitable, particularly during estrus [28,51,68]. The difference in spike-firing without accompanying changes in resting potential or membrane resistance suggests that sex differences in PV-IN intrinsic excitability may stem from voltage-gated and/or calcium-gated conductance. Furthermore, there were no sex differences in rheobase or spike-firing in PV-mGlu_5_^-/-^ mice, suggesting mGlu_5_ receptors are critical in the sex-dependent development and/or maintenance of PV-IN excitability. An important caveat is that we did not conduct simultaneous, interleaved recordings from control mice and PV-mGlu_5_^-/-^ mice, so we cannot interpret whether *Grm5* deletion makes PV-INs from female mice less excitable (or those from males more excitable) relative to controls. Nonetheless, these data suggest that basal sex differences in PV-IN excitability stem from gated ion channels or effectors regulated by mGlu_5_ receptors, potentially including calcium-activated potassium channels [47,69,70], TRP cation channels [71,72], and molecules related to Gα_q_signaling.

The current findings reveal sex differences in mGlu_1/5_ receptor signaling in PV-INs, as activation of group 1 mGlu receptors induced larger depolarizing currents in male PFC PV-INs compared to females (Supplemental Figure S6). Furthermore, the MTEP blocked DHPG-induced inward currents in both sexes but LY367385 only reached significance in male mice. These findings suggest that mGlu_1_ receptors have enhanced function in PV-INs from male mice. mGlu receptor signaling is regulated at several levels that could vary by sex. One possibility is that *Grm1* could display sex differences in splicing [73], leading to differences in subcellular protein expression and effector signaling [74,75]. Indeed, studies in hippocampal interneurons found that mGlu_1_ and mGlu_5_ receptors can activate distinct signaling pathways to regulate calcium mobilization [72], illustrating that mGlu_1_ and mGlu_5_ receptors could depolarize PV-INs through divergent mechanisms and these could also vary by sex. Finally, the propensity for mGlu_1_ and mGlu_5_ to form homodimers versus heterodimers [76,77] represents an intriguing potential source of sex differences in receptor function. Future studies should address these exciting possibilities.

In addition to membrane physiology, our findings reveal sex differences in mGlu_1_ and mGlu_5_ receptor regulation of excitatory synaptic strength on PV-INs. Previous studies found that mGlu_5_ receptors potentiate spontaneous excitatory drive onto PFC pyramidal cells in male rats [57], whereas mGlu_1_ receptors serve this role on somatostatin interneurons in both male and female mice [32,54]. Here, we found a striking sex difference in mGlu receptor subtype function on PV-INs, where mGlu_1_ enhanced excitatory drive in male mice, but mGlu_5_ receptors enhanced excitatory drive in females. To our surprise, genetically reducing mGlu_5_ receptor expression in female mice unmasked mGlu_1_ receptor-dependent responses. These findings suggest that mGlu_5_ receptors may repress mGlu_1_ receptor signaling in PV-INs of female mice, potentially at the level of gene expression [78], receptor heterodimerization [76,77], or homodimer signaling crosstalk [79]. Another possibility is that differential coupling with endocannabinoid systems could mediate sex differences in excitatory drive. Indeed, we found that mGlu_5_ receptor activation potentiated DSE in PV-INs from male but not female mice. Thus, in males, it is possible that mGlu_5_ receptor activation exerts simultaneous but nullifying actions to amplify excitatory drive and increase endocannabinoid-mediated inhibition in PV-INs. Importantly, we note these studies are limited by non-specific assessments of excitatory transmission, so future work should identify the sources of glutamate underlying these effects.

Pharmacological antagonism and genetic deletion of mGlu_5_ receptors reduce voluntary ethanol consumption, locomotor stimulation, and interoceptive effects [33,84–88]. Here, we found that genetic inhibition of mGlu_5_ receptors on PV-INs decreased binge drinking in female but not male mice. We had previously observed that IA ethanol precipitates sex-specific changes in synaptic strength on PV-INs, such that drinking reduces and enhances excitatory drive onto PV-INs in female mice but enhances sEPSC frequency onto PV-INs of male mice [28]. The current findings suggest that mGlu_5_ receptors mediate the reduction in excitatory drive onto PV-INs and this phenomenon facilitates escalated drinking in female mice. Results from our locomotor activity experiment may potentially implicate changes in ethanol’s stimulant properties. An intriguing hypothesis for future studies is that female mice escalate drinking to overcome tolerance to ethanol’s stimulant effects, and this process is regulated by mGlu_5_ signaling on PV-INs. Interestingly, previous studies using systemic pharmacology or global genetic deletion of mGlu_5_ receptors have found decreases in both drinking and stimulant and/or interoceptive effects; thus, the present findings indicate a curious cell type-specific disassociation of these attributes which merits further investigation. One important caveat to the present studies is that PV-expressing cells outside the PFC may have also contributed to the observed behavioral effects.

The current findings suggest that mGlu_5_ receptors drive sex differences in PV-IN membrane properties and synaptic plasticity. However, mGlu_5_ receptors are one of the most widely expressed GPCRs but most cell types do not express overt physiological sex differences. Even in the closely related somatostatin-expressing interneurons, we and others found minimal evidence for sex differences in basal properties [42,89–91] or in mGlu receptor signaling [29,92]. These findings beg the question, what factors make PV-INs sensitive to sex-dependent mGlu receptor signaling? Sex steroid hormones represent the most likely candidates. Clemens et al. [68] recently demonstrated that the excitability of PV-INs in somatosensory cortex is regulated by estradiol acting on membrane-associated estrogen receptor-beta (ERβ). Within neocortex, ERβ is highly localized within PV-INs [68,93], providing a compelling molecular substrate to propagate sex-dependent signaling. Across many brain regions and cell types, cell-surface ERs interact with group 1 mGlu receptors to regulate excitability and synaptic transmission [37,38,94–96]. In addition to estrogens, progesterone is another major sex hormone that can be metabolized into neurosteroids that alter GABA circuit function [97–99]. Previous studies have also linked testosterone with mGlu synaptic plasticity and eCBs [94,100]; thus, there are several hormone mechanisms that might mediate sex differences in PV-IN function.

### Conclusion

Collectively, these findings illustrate that mGlu_1_ and mGlu_5_ receptors regulate multiple aspects of PFC PV-IN physiology in a sex-dependent manner. Gaining a better understanding of the cellular and molecular mechanisms that underlie sex differences in brain function should help guide the development and translation of novel therapeutic approaches for AUD and other psychiatric diseases.

## Acknowledgments

We thank members of the University of Pittsburgh Department of Psychiatry and Translational Neuroscience Program for stimulating discussions.

## Author Contributions

CBF – Conceptualization, Investigation, Writing – Original Draft, Visualization; NDJ – Investigation; RHC – Investigation; LGC – Investigation; SMT – Investigation, Project administration; MLS – Conceptualization, Writing – Review & Editing; MEJ – Conceptualization, Investigation, Writing – Review & Editing, Supervision, Funding Acquisition.

## Supplemental Figure Legends

**Figure S1.**
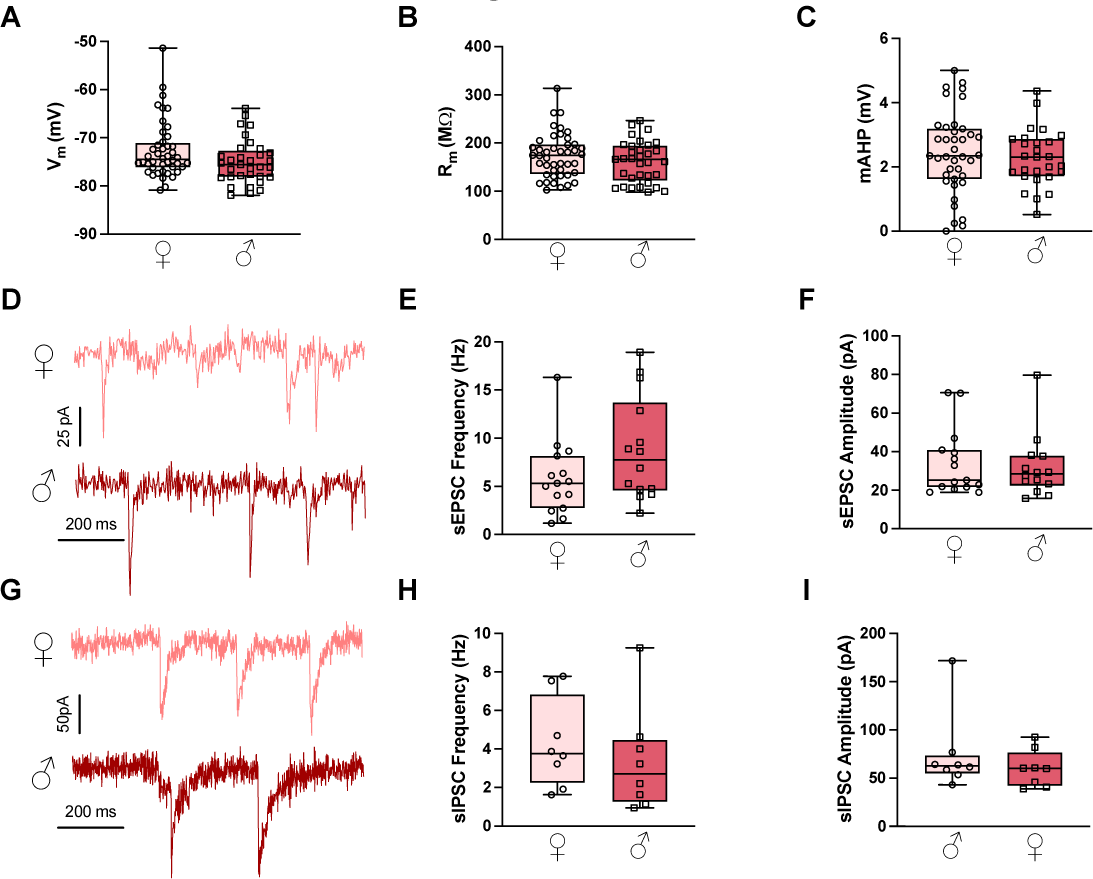
PV-INs display comparable resting membrane properties and basal synaptic strength across sexes. No sex differences in **(a)** resting membrane potential (V_m_), **(b)** membrane resistance (R_m_), or **(c)** medium afterhyperpolarization (mAHP) observed in PV-INs from male and female mice. n/N= 33-44/8 cells/mice per group. **(d)** Representative traces of baseline spontaneous excitatory postsynaptic currents (sEPSCs) from female (top, pink) and male (bottom, red) PV-INs. **(e-f)** No sex differences in basal synaptic strength onto PV-INs. n/N= 14-15/8-10 cells/mice per group. **(g)** Representative traces of baseline spontaneous inhibitory postsynaptic currents (sIPSCs) from PV-INs from female (top, pink) and male (bottom, red) mice. No sex differences in average **(h)** sIPSC frequency or **(i)** sIPSC amplitude onto PFC PV-INs. n/N= 8/3 cells/mice per group.

**Figure S2.**
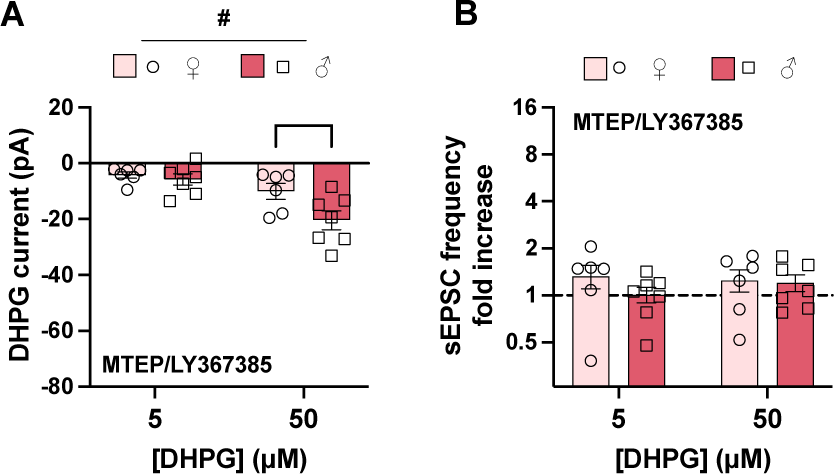
Dual application of MTEP and LY367385 attenuated DHPG currents and blocked increase in synaptic transmission in PFC PV-INs. **(a)** DHPG currents in the presence of a combination of MTEP (3 μM) and mGlu_1_ antagonist LY367385 (100 μM). Currents are reduced compared to control or single NAM experiments (Figure 1); however, DHPG still exerted a concentration-dependent and sex-dependent depolarizing current (Two-way ANOVA, main effect of concentration: F_1,11_ = 20.1, p<0.001, main effect of sex: F_1,11_ = 4.5, p<0.06, concentration x sex interaction: F_1,11_ = 3.7, p<0.08, *:p<0.05, **:p<0.01 Bonferonni post-test). **(b)** MTEP/LY367385 application blocked the ability of DHPG to enhance excitatory drive onto PFC PV-INs (Two-way ANOVA, main effect of concentration: F_1,11_ = 0.3, n.s.; one-sample t-test t_25_ = 1.2, n.s.). n/N = 6-7/3.

**Figure S3.**
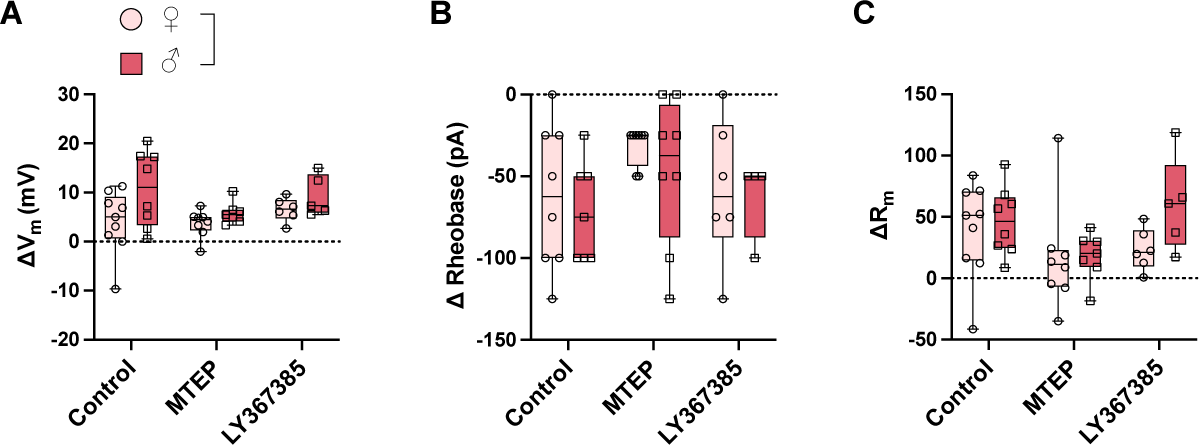
Male PV-INs undergo greater depolarization following mGlu_1_ and mGlu_5_ receptor stimulation. Membrane properties of PFC PV-INs assessed under current clamp configuration during end of 50 μM DHPG wash-on. Change calculated as the difference relative to the baseline before DHPG wash-on. **(a)** PV-INs from males exhibit a greater DHPG-induced depolarization in resting membrane potential (Vm) than cells from females (Two-way ANOVA, main effect of sex: F_2,37_ = 1.6, *:p<0.05). No main effect of sex or antagonist on the change in **(b)** rheobase or **(c)** membrane resistance (R_m_) in PV-INs. n/N = 6-9/8-10 cells/mice per group.

**Figure S4.**
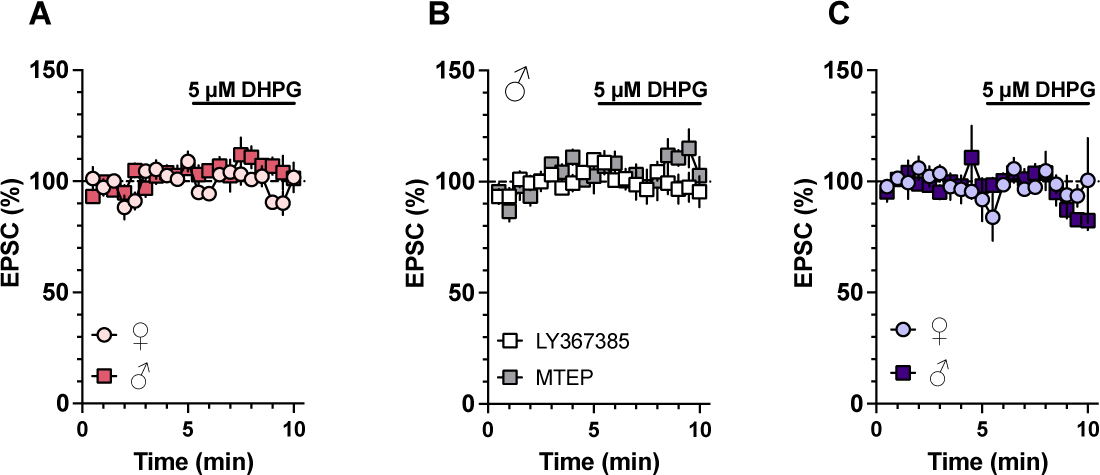
Threshold DHPG (5 μM) did not affect the amplitude of evoked EPSCs on PV-INs. Prior to depolarization-induced suppression of inhibition experiments, threshold DHPG (5 μM) was applied to cells for 5 minutes while electrically evoked EPSCs were monitored. Threshold DHPG had no effect on EPSC amplitude in control recordings **(a)**, in the presence of LY367385 or MTEP **(b)**, or in PV-mGlu_5_^-/-^ mice **(c)**. n/N = 5-8/4-5 mice per group.

**Figure S5.**
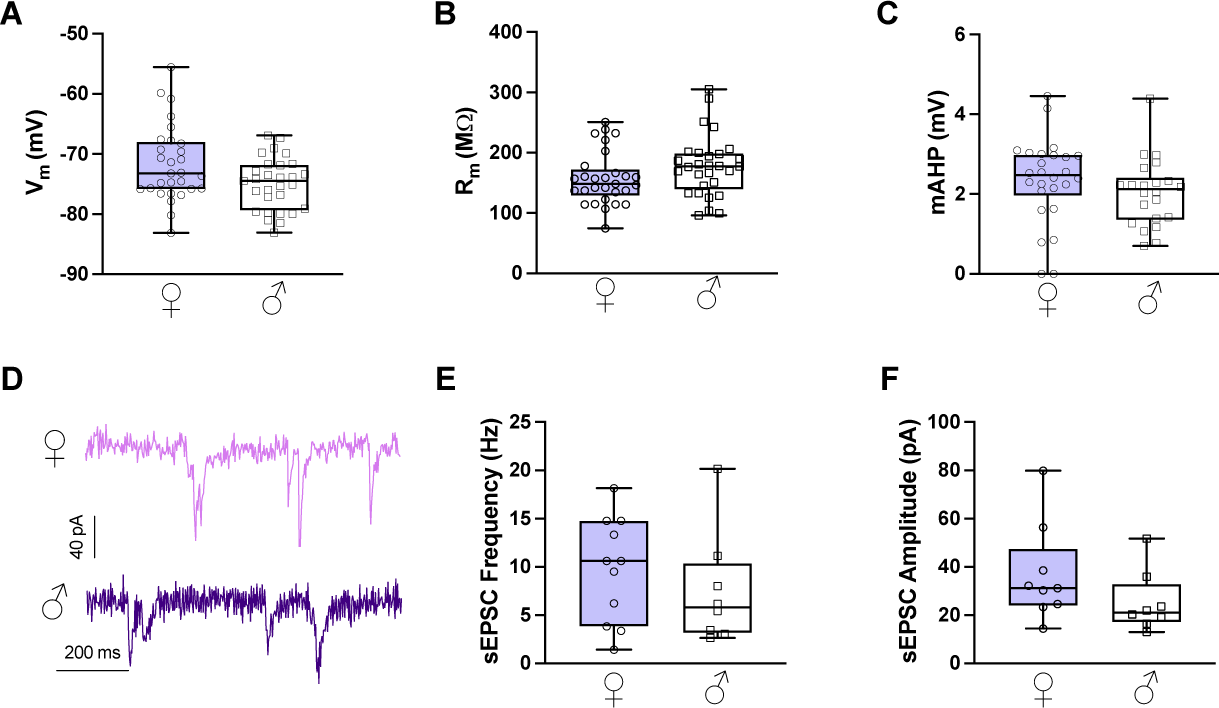
PV-mGlu_5_^-/-^ mice display comparable basal membrane properties and synaptic strength across sexes. PV-mGlu_5_^-/-^ mice display no sex differences in **(a)** resting membrane potential (V_m_), **(b)** membrane resistance (R_m_), or **(c)** medium afterhyperpolarization (mAHP). **(d)** Representative traces of baseline EPSCs from PV-mGlu_5_^-/-^ female (top, light purple) and male (bottom, dark purple) mice. n/N= 22-24/8-10 cells/mice per group. **(e-f)** PV-mGlu_5_^-/-^ display no sex differences in basal synaptic strength onto PV-INs. n/N= 8-9/3-4 cells/mice per group.

**Figure S6.**
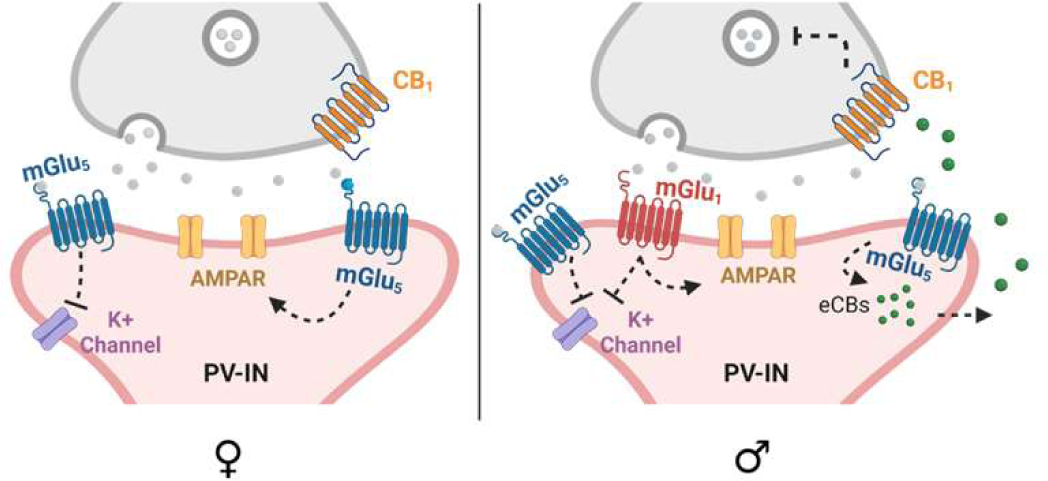
Schematic depicting proposed sex differences in mGlu_1_ and mGlu_5_ receptor signaling in PFC PV-INs. In female mice, mGlu_5_ receptor signaling depolarizes PV-INs, likely through inhibition of potassium channels. In males, cell significant depolarization of PV-INs is mediated by both mGlu_1_ and mGlu_5_ receptor signaling. Synaptic strength onto PV-INs is enhanced by mGlu_5_ receptor activation in female mice; by contrast, in males synaptic strength onto PV-INs is potentiated by mGlu_1_ receptor activation. Finally, mGlu_5_ receptor activation potentiates endocannabinoid short-term plasticity onto PV-INs from male mice but not females.

