## Supplemental Tables for "Parvalbumin interneuron mGlu_5_ receptors govern sex differences in prefrontal cortex physiology and binge drinking"

Supplemental Table 1. Means ± SEM for key findings

| Figure 1 | Sex | Condition | Mean | SEM | | n cells |
|---|---|---|---|---|---|---|
| 1C | Female | Baseline | 150.0 | 10.5 | pA | 44 |
| | Male | Baseline | 191.1 | 12.8 | | 31 |
| 1E | Female | Control | -11.3 | 1.9 | pA | 11 |
| | Female | MTEP | -5.8 | 1.4 | | 11 |
| | Female | LY367385 | -6.6 | 1.0 | | 10 |
| | Male | Control | -14.7 | 3.0 | | 10 |
| | Male | MTEP | -11.6 | 3.1 | | 9 |
| | Male | LY367385 | -8.3 | 3.4 | | 11 |
| 1F | Female | Control | -32.2 | 3.9 | pA | 14 |
| | Female | MTEP | -16.6 | 4.3 | | 11 |
| | Female | LY367385 | -25.9 | 4.3 | | 9 |
| | Male | Control | -50.4 | 6.4 | | 13 |
| | Male | MTEP | -27.2 | 5.1 | | 8 |
| | Male | LY367385 | -23.8 | 5.3 | | 8 |
| **Figure 2** | **Sex** | **Condition** | **Mean** | **SEM** | | **n cells** |
| 2B | Female | Control | 1.4 | 0.2 | fold | 12 |
| | Female | MTEP | 1.1 | 0.1 | change | 11 |
| | Female | LY367385 | 1.2 | 0.2 | | 8 |
| | Male | Control | 1.7 | 0.4 | | 10 |
| | Male | MTEP | 1.3 | 0.2 | | 9 |
| | Male | LY367385 | 0.8 | 0.1 | | 11 |
| 2C | Female | Control | 3.6 | 0.6 | fold | 15 |
| | Female | MTEP | 2.0 | 0.5 | change | 11 |
| | Female | LY367385 | 3.1 | 0.8 | | 7 |
| | Male | Control | 2.7 | 0.5 | | 13 |
| | Male | MTEP | 2.2 | 0.5 | | 8 |
| | Male | LY367385 | 1.1 | 0.2 | | 7 |
| 2F | Female | Baseline | 79.8 | 5.7 | EPSC | 8 |
| | Female | DHPG | 84.0 | 6.2 | (%) | 5 |
| | Male | Baseline | 81.9 | 3.9 | | 14 |
| | Male | DHPG | 63.2 | 4.8 | | 9 |
| 2I | LY367385 | Baseline | 79.6 | 5.1 | EPSC | 9 |
| | LY367385 | DHPG | 60.9 | 5.3 | (%) | 7 |
| | MTEP | Baseline | 74.9 | 4.6 | | 6 |
| | MTEP | DHPG | 76.3 | 5.3 | | 6 |
| **Figure 3** | **Sex** | **Condition** | **Mean** | **SEM** | | **n cells** |
| 3C | Female | Baseline | 151.6 | 13.1 | pA | 32 |
| | Male | Baseline | 158.3 | 12.2 | | 30 |
| 3E | Female | Control | -11.9 | 1.7 | pA | 7 |
| | Female | MTEP | -4.3 | 2.2 | | 8 |
| | Female | LY367385 | -3.9 | 3.8 | | 7 |
| | Male | Control | -5.2 | 4.5 | | 8 |
| | Male | MTEP | -3.2 | 2.2 | | 7 |
| | Male | LY367385 | -6.4 | 2.3 | | 8 |

| | | | | | | |
|---|---|---|---|---|---|---|
| **3F** | Female | Control | -26.9 | 5.5 | pA | 8 |
| | Female | MTEP | -11.0 | 4.5 | | 7 |
| | Female | LY367385 | -3.6 | 8.6 | | 5 |
| | Male | Control | -25.2 | 4.7 | | 7 |
| | Male | MTEP | -14.3 | 2.9 | | 5 |
| | Male | LY367385 | -16.7 | 5.0 | | 8 |
| **Figure 4** | **Sex** | **Condition** | **Mean** | **SEM** | **Unit** | **n cells** |
| **4B** | Female | Control | 1.2 | 0.1 | fold change | 9 |
| | Female | MTEP | 1.2 | 0.2 | | 8 |
| | Female | LY367385 | 1.1 | 0.2 | | 7 |
| | Male | Control | 1.3 | 0.2 | | 8 |
| | Male | MTEP | 1.3 | 0.1 | | 7 |
| | Male | LY367385 | 1.0 | 0.1 | | 8 |
| **4C** | Female | Control | 2.6 | 0.4 | fold change | 9 |
| | Female | MTEP | 2.3 | 0.8 | | 7 |
| | Female | LY367385 | 1.4 | 0.3 | | 5 |
| | Male | Control | 4.2 | 0.8 | | 7 |
| | Male | MTEP | 2.2 | 0.6 | | 5 |
| | Male | LY367385 | 1.8 | 0.2 | | 8 |
| **Figure 5** | **Sex** | **Genotype** | **Mean** | **SEM** | **Unit** | **N mice** |
| **5B** | Female | WT | 19.3 | 1.0 | g/kg/day | 12 |
| | Female | PV-mGlu$_5^{-/-}$ | 14.7 | 1.5 | | 6 |
| | Male | WT | 12.3 | 1.4 | | 8 |
| | Male | PV-mGlu$_5^{-/-}$ | 10.9 | 0.9 | | 12 |
| **5D** | Female | WT | 11.5 | 2.1 | distance (m) | 7 |
| | Female | PV-mGlu$_5^{-/-}$ | 26.7 | 4.0 | | 8 |
| | Male | WT | 10.4 | 2.5 | | 7 |
| | Male | PV-mGlu$_5^{-/-}$ | 24.0 | 5.3 | | 7 |
| **5E** | Female | WT | 8.0 | 1.4 | distance (m) | 7 |
| | Female | PV-mGlu$_5^{-/-}$ | 12.1 | 1.8 | | 8 |
| | Male | WT | 5.7 | 1.2 | | 7 |
| | Male | PV-mGlu$_5^{-/-}$ | 12.7 | 2.5 | | 7 |

Supplemental Table 2. Three-way ANOVA main effects and interactions

| **Three-way ANOVA: 5μM and 50μM DHPG holding current PV** | | | | |
|---|---|---|---|---|
| **1E-1F** | **Fixed effects (type III)** | **P value** | **Summary** | **(P < 0.05)?** |
| | antagonists | 0.0006 | *** | Yes |
| | DHPG concentration | <0.0001 | **** | Yes |
| | sex | 0.03 | * | Yes |
| | antagonists x DHPG concentration | 0.0062 | ** | Yes |
| | antagonists x sex | 0.2567 | ns | No |
| | DHPG concentration x sex | 0.1003 | ns | No |
| | antagonists x DHPG concentration x sex | 0.0752 | ns | No |
| **Three-way ANOVA: 5μM and 50μM DHPG sEPSCs PV** | | | | |
| **2B-2C** | **Fixed effects (type III)** | **P value** | **Summary** | **(P < 0.05)?** |
| | antagonists | 0.0632 | ns | No |
| | DHPG concentration | <0.0001 | **** | Yes |
| | sex | 0.175 | ns | No |
| | antagonists x DHPG concentration | 0.3569 | ns | No |
| | antagonists x sex | 0.2121 | ns | No |
| | DHPG concentration x sex | 0.0454 | * | Yes |
| | antagonists x DHPG concentration x sex | 0.2395 | ns | No |
| **Three-way ANOVA: 5μM and 50μM DHPG holding current PV-mGlu5-/-** | | | | |
| **3E-3F** | **Fixed effects (type III)** | **P value** | **Summary** | **(P < 0.05)?** |
| | antagonists | 0.0717 | ns | No |
| | DHPG concentration | <0.0001 | **** | Yes |
| | sex | 0.7383 | ns | No |
| | antagonists x DHPG concentration | 0.0128 | * | Yes |
| | antagonists x sex | 0.3509 | ns | No |
| | DHPG concentration x sex | 0.2061 | ns | No |
| | antagonists x DHPG concentration x sex | 0.4975 | ns | No |
| **Three-way ANOVA: 5μM and 50μM DHPG sEPSCs PV-mGlu5-/-** | | | | |
| **4B-4C** | **Fixed effects (type III)** | **P value** | **Summary** | **(P < 0.05)?** |
| | antagonists | 0.0051 | ** | Yes |
| | DHPG concentration | <0.0001 | **** | Yes |
| | sex | 0.2099 | ns | No |
| | antagonists x DHPG concentration | 0.0071 | ** | Yes |
| | antagonists x sex | 0.3206 | ns | No |
| | DHPG concentration x sex | 0.1296 | ns | No |
| | antagonists x DHPG concentration x sex | 0.2682 | ns | No |

| F (DFn, DFd) |
|---|
| F (2, 62) = 8.373 |
| F (1, 51) = 143.5 |
| F (1, 62) = 4.936 |
| F (2, 51) = 5.626 |
| F (2, 62) = 1.390 |
| F (1, 51) = 2.801 |
| F (2, 51) = 2.723 |

| F (DFn, DFd) |
|---|
| F (2, 61) = 2.890 |
| F (1, 49) = 36.87 |
| F (1, 61) = 1.883 |
| F (2, 49) = 1.052 |
| F (2, 61) = 1.591 |
| F (1, 49) = 4.215 |
| F (2, 49) = 1.472 |

| F (DFn, DFd) |
|---|
| F (2, 40) = 2.817 |
| F (1, 33) = 47.65 |
| F (1, 40) = 0.1132 |
| F (2, 33) = 4.993 |
| F (2, 40) = 1.075 |
| F (1, 33) = 1.664 |
| F (2, 33) = 0.7132 |

| F (DFn, DFd) |
|---|
| F (2, 41) = 6.011 |
| F (1, 35) = 38.96 |
| F (1, 41) = 1.622 |
| F (2, 35) = 5.720 |
| F (2, 41) = 1.170 |
| F (1, 35) = 2.409 |
| F (2, 35) = 1.367 |
